# Sarek: A portable workflow for whole-genome sequencing analysis of germline and somatic variants

**DOI:** 10.1101/316976

**Authors:** Maxime Garcia, Szilveszter Juhos, Malin Larsson, Pall I. Olason, Marcel Martin, Jesper Eisfeldt, Sebastian DiLorenzo, Johanna Sandgren, Teresita Diaz de Ståhl, Valtteri Wirta, Monica Nistér, Björn Nystedt, Max Käller

## Abstract

**Summary:** Whole-genome sequencing (WGS) is a cornerstone of precision medicine, but portable and reproducible open-source workflows for WGS analyses of germline and somatic variants are lacking. We present Sarek, a modular, comprehensive, and easy-to-install workflow, combining a range of software for the identification and annotation of single-nucleotide variants (SNVs), insertion and deletion variants (indels), structural variants, tumor sample heterogeneity, and karyotyping from germline or paired tumor/normal samples. Sarek is implemented in a bioinformatics workflow language (Nextflow) with Docker and Singularity compatible containers, ensuring easy deployment and full reproducibility at any Linux based compute cluster or cloud computing environment. Sarek supports the human reference genomes GRCh37 and GRCh38, and can readily be used both as a core production workflow at sequencing facilities and as a powerful stand-alone tool for individual research groups.

**Availability:** Source code and instructions for local installation are available at GitHub (https://github.com/SciLifeLab/Sarek) under the MIT open-source license, and we invite the research community to contribute additional functionality as a collaborative open-source development project.

## 1. Introduction

Whole-genome sequencing (WGS) opens up new avenues for research and diagnostics, with many large national and international initiatives already launched worldwide^1–6^. While many sequencing facilities provide WGS germline and somatic variant calling as a part of their service, these workflows are typically difficult to deploy elsewhere, limiting transparency, reproducibility, and re-usability. Sarek offers a robust and portable analysis workflow, handling both germline and somatic variant detection and annotation from WGS data, including a range of the currently most widely used software and data resources in the field (Fig 1A, Supplementary Information S1). This is of particular importance for somatic variant calling, where a combination of tools is required to achieve optimal sensitivity and specificity^7^. The workflow is easy to install, and runs on any Linux based computers, high-performance compute clusters, and cloud solutions. Sarek supports both GRCh37 and GRCh38, and much of its features are also applicable to whole-exome sequencing (WES) and other targeted sequencing approaches. Here we present the design, usage and performance of Sarek, including resource utilization and qualitative benchmarking on somatic variant calling from *in silico* and real datasets.

**Figure 1.**
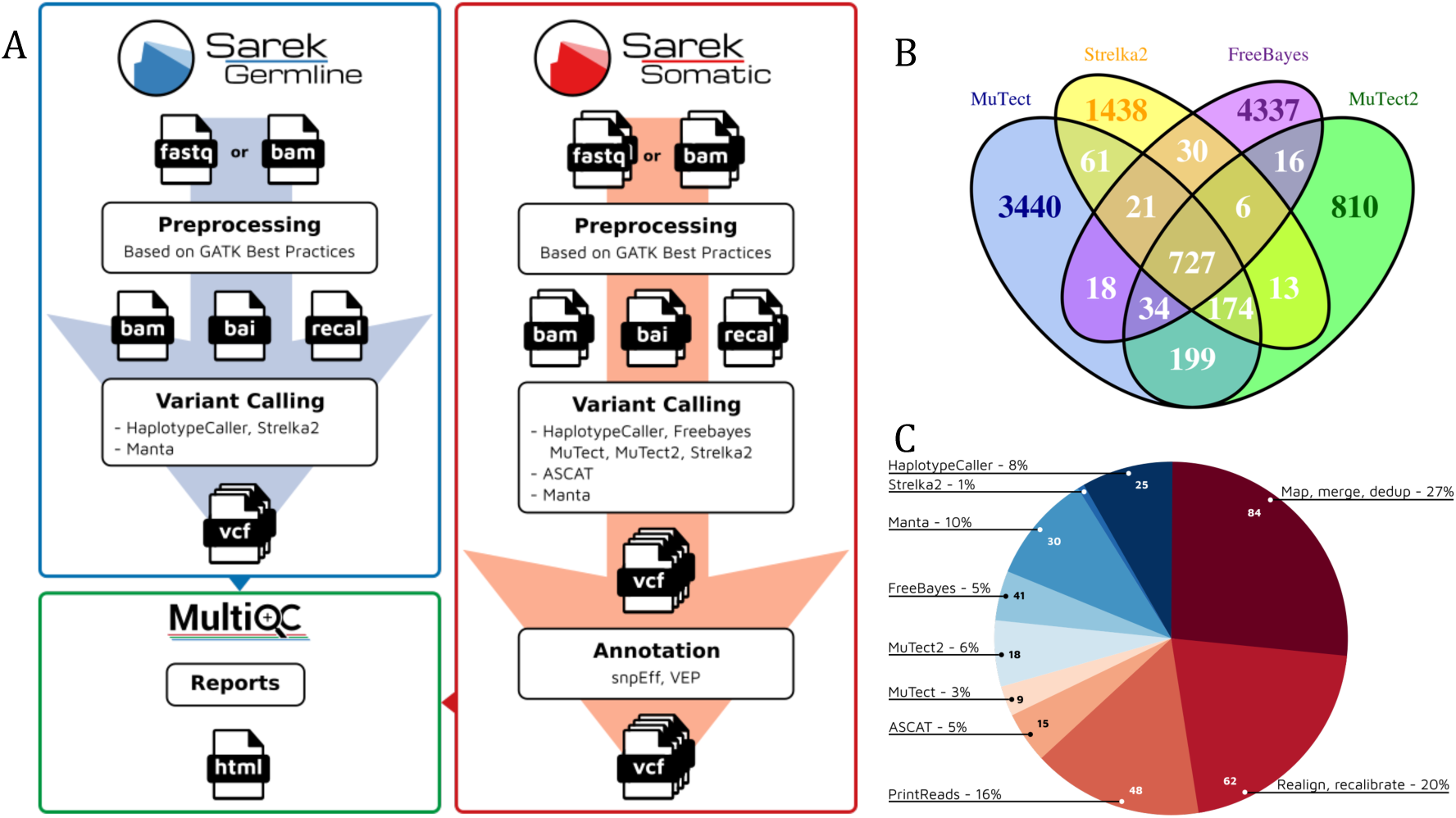
(A) Schematic overview of the Sarek workflows for germline and somatic variants. (B) Venn diagram showing the number and overlap of somatic single nucleotide variants (SNVs) detected by MuTect, MuTect2, Strelka2 and FreeBayes from a whole-genome sequencing (WGS) medulloblastoma dataset^7^. (C) Wall-clock time in hours for a complete Sarek run on a 90X/90X sequence coverage tumor/normal sample on a 16-core node.

## 2. Implementation

### Germline and somatic variant detection and annotation

Sarek is implemented in the Nextflow workflow language^8^ to ensure portability, robustness, and efficient utilization of computer resources. Sarek is open source and has containers compatible with Docker (http://www.docker.com) and Singularity^9^ for easy installation and full reproducibility. In the pre-processing step, sequence reads are aligned to the reference genome with BWA-MEM^10^, followed by realignment and recalibration with GATK^11^. Germline mutations are detected from a single sample both with HaplotypeCaller11 according to the GATK best-practice recommendations, and with Strelka2^12^, with structural variations detected with Manta^13^. After joint realignment of the sequence reads from tumor and normal samples from the same individual, somatic mutations are identified using a range of somatic variant callers, including MuTect^14^, MuTect2^14^, Strelka2^12^, FreeBayes^15^, Manta^12^, and ASCAT^16^, covering the detection of SNVs, indels, and structural variants (including copy-number variation), as well as karyotype and sample purity. Additional variant callers can easily be incorporated, further extending the capability of Sarek. The resulting variants from each variant caller are annotated for potential functional effects with snpEff ^17^ and VEP^18^.

### Sample quality control

During the run of the workflow, quality control (QC) processes are launched to scan the result files, which can be imported into MultiQC^19^ for an overview of quality control metrics. MultiQC modules are available to present relevant statistics and plots for the input FASTQ files by FastQC (http://www.bioinformatics.babraham.ac.uk/projects/fastqc), alignment and coverage descriptors by BamQC (https://github.com/s-andrews/BamQC), QualiMap^20^, BCFtools^21^ and Samtools^22^, MarkDuplicates statistics using Picard (http://broadinstitute.github.io/picard/), germline and somatic variant analysis results by VCFtools^23^ and snpEff^17^. MultiQC aggregates these reports into a single HTML review per sample for easy visualisation.

### Variant detection

We performed somatic variant calling with Sarek using a medulloblastoma tumor/normal pair dataset from the ICGC PedBrain Tumor project, with ~90X short-read sequencing coverage each for the normal and tumor samples (“MB90”, accession number EGAD00001001859). For this sample, a "Gold Set" of somatic variants has been released by the ICGC based on very deep (>300X) WGS sequencing^7^. In addition, we also analysed two *in silico* datasets from the DREAM challenge^24^ (“IS2”, accession number SRX1025978, “IS3” accession number SRX1026041), modelling cancer samples of varying complexity, with 40X sequencing coverage each for the normal and tumor samples. By combining the output from four variant calling softwares (MuTect, MuTect2, Strelka2, FreeBayes) (Fig. 1B), Sarek provided robust recall and precision performance across the three datasets well comparable to the results from previous benchmarking studies^7,24^ for both SNVs, indels and structural variants (Table 1, Supplementary Information S2).

**Table 1.** Example recall, precision, and F-score values based on the combined output of multiple somatic variant callers for SNVs, indels and structural variants produced in a Sarek run for two *in silico* (IS2, IS3) and one real (MB90) dataset. The number of variants of each type in the True Set is indicated in brackets for each dataset.

| SNVs detected by at least 3 out of 4 callers | Recall | Precision | F-score |
| --- | --- | --- | --- |
| IS2 (n=4332) | 0.97 | 0.74 | 0.84 |
| IS3 (n=7903) | 0.90 | 0.98 | 0.94 |
| MB90 (n=1286) | 0.70 | 0.93 | 0.80 |
| Indels detected by at least 2 out of 3 callers |  |  |  |
| IS2 (n=0)* | NA | NA | NA |
| IS3 (n=8018) | 0.61 | 0.96 | 0.75 |
| MB90 (n=351) | 0.60 | 0.55 | 0.57 |
| Structural variants detected by Manta |  |  |  |
| IS2 (n=655) | 0.70 | 0.86 | 0.77 |
| IS3 (n=2886) | 0.68 | 0.85 | 0.76 |
| MB90** | NA | NA | NA |

\* No true indels are present in the *in silico* IS2 dataset
\*\* No True Set is available for structural variants in the MB90 sample

### Resource usage

To test speed and resource usage, Sarek was run in somatic mode on the MB90 dataset on an HPC cluster (250 dual CPU Intel Xeon E5-2630 v3, 2.40 GHz; 256 GB, 4 TB storage, and 16 cores per node; 1.1 PB Network-attached storage (NAS) connected via Gigabit Ethernet and 4xQDR Infiniband), with the SLURM workload manager. Computations were done on a single node with intermediate files stored on the local scratch area and final files copied to the NAS. A full run starting from FASTQ files including all implemented somatic variant callers and annotation steps required about 14 days on a 16-core node, with a total storage output about four times larger than the original input data (Fig. 1C, Supplementary Information S3). We note that a significant fraction of the computational and storage needs are attributed to the realignment step, which can be omitted if MuTect is excluded, since this step is integrated in the other variant calling softwares.

### Installation and usage

Operating from a computer system with a local installation of Nextflow and support for Docker or Singularity containers, Nextflow can automatically fetch the Sarek source tree from GitHub. Notably, Sarek comes with pre-built containers for all its dependency tools, thus avoiding cumbersome local software installations for the end user. A full workflow starting from FASTQ files including mapping, realignment, recalibration, variant calling, and annotation can be invoked as below

~~~
> nextflow run SciLifeLab/Sarek/main.nf --sample samples.tsv --step mapping
> nextflow run SciLifeLab/Sarek/somatic.nf --tools mutect2,strelka,freebayes,manta,ascat
> nextflow run SciLifeLab/Sarek/annotate.nf --tools snpEff,VEP
> nextflow run SciLifeLab/Sarek/runMultiQC.nf
~~~

A number of configuration files included in the installation allows adjustment of software parameters and tailoring of the workflow to specific user needs. Incomplete runs are easily restarted from the point of failure, by simply invoking the same command as above, as the workflow will not recreate already existing output files unless forced to do so. To verify the installation, the workflow comes with a small test dataset using a part of GRCh37 as the reference (https://github.com/SciLifeLab/Sarek-data).

## 3. Conclusions

Sarek is a portable and reproducible workflow to detect germline and somatic variants from WGS data. Sarek has recently been implemented in routine production at the National Genomics Infrastructure at SciLifeLab (www.scilifelab.se), one of the largest sequencing facilities in Europe, and has also been successfully tested as a stand-alone tool by several clinical research groups. A wide range of software is already included in Sarek, and we expect the ongoing implementation of GATK4^11^ and support for CRAM formats^25^ to enhance the speed and reduce storage needs. Downstream ranking and visualization modules operating on the Sarek output are already under development, with the aim to support clinical decisions in health care.

## Acknowledgements

We are grateful for valuable input from the Oslo University Hospital bioinformatics core facility (Oslo university hospital), the T Martinsson lab (Gothenburg university), and the A-C Syvänen lab (Uppsala university). The computations were performed on resources provided by the National Genomics Infrastructure (NGI) and Uppsala Multidisciplinary Center for Advanced Computational Science (UPPMAX). We thank Dr. Jonas Söderberg for help with graphical design.

